# ImpRes: A robust FRAP framework to quantify fast diffusion of cytoplasmic probes

**DOI:** 10.64898/2026.08.14.744877

**Authors:** Olivier Destrian, René-Marc Mège, Benoît Goyeau, Morgan Chabanon

## Abstract

Diffusion within the cytoplasm is fundamental to numerous biological processes. Fluorescence recovery after photobleaching (FRAP) is one of the most common method for quantifying molecular diffusivity in living cells using standard laser scanning confocal microscopy (LSCM). However, accurately measuring fast cytoplasmic diffusion (typically >10 µm^2^/s) is challenging due to rapid recovery kinetics, weak signal-to-noise ratios, post-bleach signal artifacts, and spatial restrictions affecting normalization. While individual challenges have been addressed in specific contexts, a simple and robust framework to quantify cytoplasmic diffusivity remains elusive. Here, we present a FRAP methodology specifically designed to overcome these obstacles. By utilizing the Gaussian function – the impulse response (ImpRes) of the diffusion equation in an infinite medium – our approach leverages the full spatiotemporal dataset through a single-equation three-parameter fitting procedure, thus releasing restrictions to small regions of interest and arbitrary initial time-points. The methodology was validated on three datasets of increasing complexity: *in silico* simulated recovery profiles, *in vitro* data from FITC-dextran in glycerol solution, and live-cell imaging of cytoplasmic free-GFP. Systematic comparison with existing models demonstrates that the ImpRes approach significantly reduces sensitivity to noise and imperfect fluorescence normalization, while remaining robust against short-term biases, such as transient probe photo-activation. Given its robustness under realistic experimental conditions and its ease of implementation, the proposed FRAP methodology provides a reliable tool for quantitative cytoplasmic analysis.

## 1 Introduction

The impact of cytoplasmic crowding and structural heterogeneity on cellular functionality has gained significant interest in recent years [1, 2, 3, 4, 5]. Crowding notably constrains the intracellular diffusion of biomolecules, thereby influencing a wide array of essential biological processes, including organelle biogenesis [6], cytoskeletal dynamics [7, 8], and the orchestration of apoptosis [9]. These findings have relied on the study of intracellular molecular transport, which has been driven largely by the development of fluorescent labeling and advanced microscopy techniques, most notably fluorescence recovery after photobleaching (FRAP) [10, 11, 12].

The fundamental principle of FRAP involves the application of a high-intensity laser over a short period of time to a localized region, inducing irreversible photobleaching of fluorescent molecules. The subsequent recovery of fluorescence in the bleached area is then monitored over time. By applying appropriate mathematical models to the recovery kinetics, one can quantify molecular transport properties, most notably the effective diffusion coefficient.

Complementary techniques such as single-particle tracking (SPT) [13] and fluorescence correlation spectroscopy (FCS) [14] can provide high-precision data at the sub-micron scale, such as individual particle trajectories, residence times, and specific motility mechanisms [3, 1, 15]. However FRAP remains widely used due to its ease of interpretation for broad range of fluorescent probes and its relatively simple implementation with standard laser scanning confocal microscopy (LSCM) setups. Furthermore, FRAP is uniquely suited for rapidly obtaining large-scale, ensemble-averaged transport properties, such as the effective diffusivity of a probe within a micrometer-sized volume.

However, while most current FRAP methods are easily applicable to slowly diffusing or membranebound molecules [16], quantitative measurements of cytoplasmic diffusivity for highly mobile molecules remain challenged due to:

- Rapid recovery kinetics: For highly mobile species, the recovery phase often occurs in less than one second. This requires to scan at high-speed and several times the region of interest. As a consequence a fundamental trade-off emerges: if the laser power is too strong it bleaches the whole zone – including the reference fluorescence region [14] – but if it is too weak the signal to noise ratio (SNR) becomes unacceptable.
- Non-instantaneous photobleaching: The photobleaching process itself requires a finite amount of time, sometimes more than 100ms. When the recovery duration is itself occurring faster than 1000ms, the bleaching phase can no longer be considered instantaneous. This leads to a reduced bleaching depth and a larger blurred spatial bleaching profile, further reducing SNR and restricting the applicability of many standard models that rely on a precise, predefined bleaching geometry [17].
- Cytoplasmic heterogeneity: Standard FRAP models typically assume a homogeneous initial probe density. However, the cytoplasm is inherently heterogeneous due to the presence of organelles, cytoskeletal networks, and macromolecular crowding [3, 4]. These structures which appear as dark occlusions in the image are often mobile, creating complex, time-dependent noise patterns in the fluorescence signal that acts as an extra layer of noise to the FRAP experiments.
- Other effects that can induce biases at different time scales include transverse diffusion for samples thicker than the LSCM bleaching point spread function (PSF) [18], spatial variations in cytoplasmic thickness [19], as well as probe specific dynamics such as GFP photoconversion and photoactivation [20, 21, 22].

Given these complexities, FRAP experiments are sometimes conducted without sufficient consideration of these limitations, potentially leading to results misinterpretation. While various models have been proposed to address specific issues [23, 19, 14, 16, 24, 18, 17, 25], a comprehensive and simple methodology applicable to standard LSCM that addresses these limitations simultaneously is still lacking.

In this work, we propose a FRAP protocol and diffusivity inference model designed to alleviate these obstacles when studying fast cytoplasmic diffusion using a standard LSCM. The experimental protocol consists in generating a Gaussian bleaching profile from a local point exposure to minimize bleaching phase duration. Imaging is performed on an elongated rectangle during recovery to maximize acquisition frequency, while providing both the full spatial extent of the bleaching profile near the rectangle center and a robust fluorescence reference at the rectangle extremities. The associated FRAP model is based on the impulse response of the diffusion equation in an infinite medium, *i.e.* a Gaussian function. By leveraging the entire spatiotemporal data set in a single three-parameter fitting procedure, our methodology is not restricted to small regions of interest, and avoids biases due the choice of an arbitrary initial time-point. We test, compare, and validate this model against widely-used existing methods [16, 24] applied to three FRAP data-sets: *in silico* generated diffusion profiles with fluorescence normalization and controlled noise intensity, *in vitro* experiments with FITC-dextran in glycerol-water solutions, and *in vivio* data from live MDCK cells expressing free-GFP.

Results show that our approach significantly reduces the sensitivity to noise, imperfect fluorescence normalization, and short-term biases such as fluorescent probe photoactivation following the bleaching phase which may result in stringent fitting timerange constraints. While existing models perform adequately under ideal conditions, under realistic experimental conditions our models provides enhanced robustness without adding complexity to the fit procedure.

## 2 Materials and Methods

### 2.1 Imaging and FRAP protocol

#### 2.1.1 Microscope setup

Experiments were performed on a Zeiss LSM980-Airyscan 2 laser scanning confocal microscope (ZenBlue software) with Plan Apochromat 63X/1.4 NA oil DIC objective. A 488 nm laser line (maximum output power measured at 5mW upstream of the objective) was used for all acquisitions. The incubated enclosure was set at 37*^◦^*C and 5% CO2 for live-cell experiments and left at room temperature for in-vitro experiments. FRAP imaging phases were acquired in confocal mode with pinhole set at 1 AU. Z-stacks were captured in Airyscan multiplex SR4Y mode and processed via ZenBlue 2D Airyscan post-processing following daily detector calibration.

#### 2.1.2 FRAP experimental protocol

FRAP was performed using a combination of 0D spot bleaching and 2D scanning for imaging.

Photobleaching was performed in 0D spot mode using 100% laser power for a duration of 25 ms for live-cell experiments and between 5 ms and 15 ms for in-vitro experiments. This approach ensured rapid and sufficient bleaching depth typically ranging from 25% to 50% of total fluorescence level. The resulting bleached profile was determined by the microscope’s bleaching point spread function (PSF) and free-GFP diffusion during bleaching, typically yielding a Gaussian profile with radius of 3–6 µm in the initial post-bleach frames.

Pre-bleach and post-bleach imaging phases were captured in 2D confocal scanning mode with a low laser power around 0.1-1% depending on sample fluorescence level. To obtain high enough temporal resolution to have several images of the recovery even for fast diffusing probes such as GFP (*τ ≤* 500 ms), a narrow rectangular region of interest (ROI) was imaged (Figure 1A). The ROI width was set to 1 µm for all experiments (Figure 1B), so as to be several times smaller than the initial bleaching radius. The ROI length was set to 25 µm for live-cell experiments and between 30 µm and 40 µm for in-vitro experiments, so as to be several times larger than the initial bleaching radius. Imaged region acquisition rate was set at 50 Hz for live-cell experiments and between 15 Hz and 50 Hz for in-vitro experiments.

**Figure 1:**
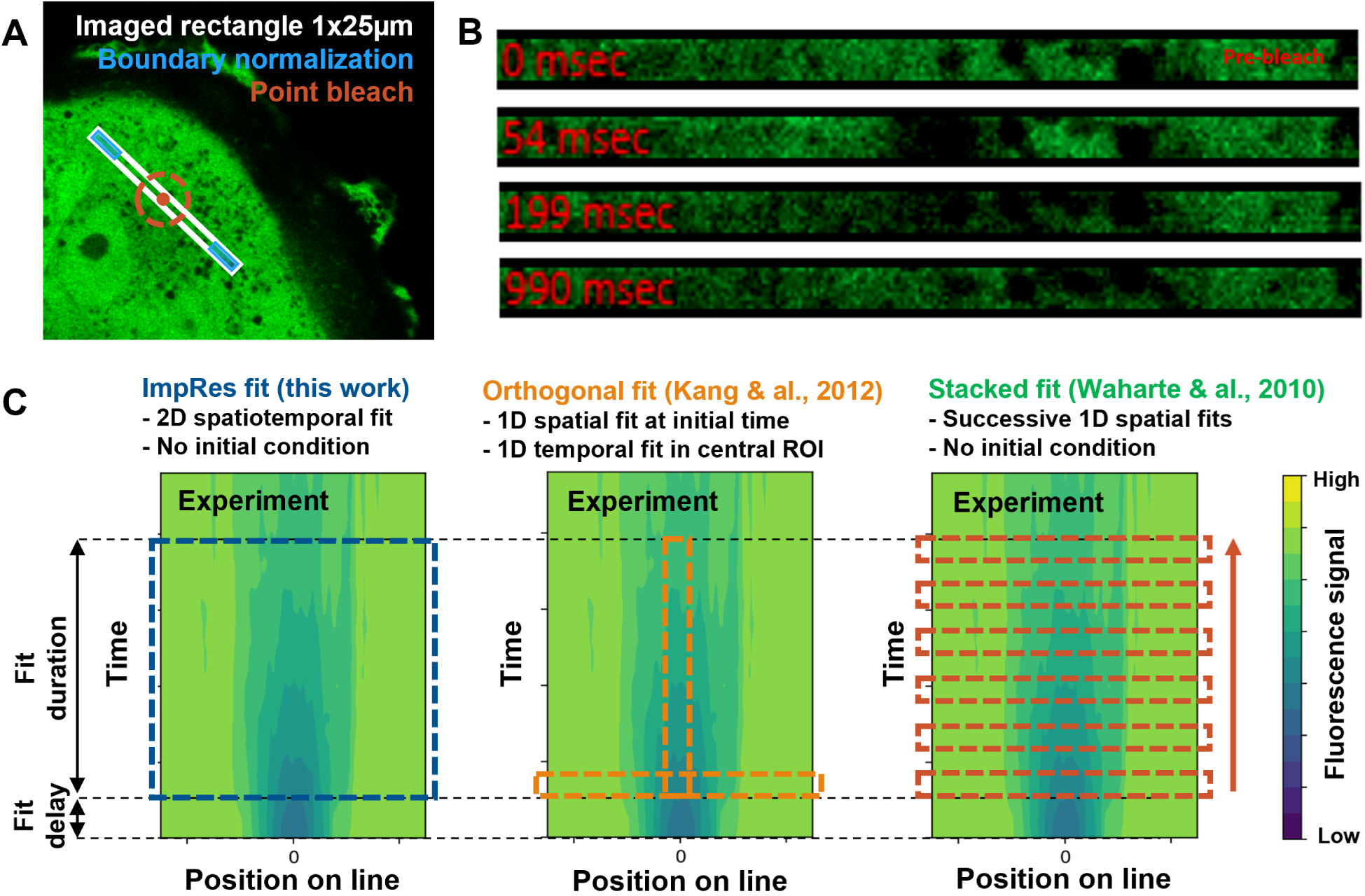
FRAP methodological approach and comparison (A): Fluorescence confocal image of a MDCK cell expressing free-GFP. Fluorescence signal is recorded on a 25×1 µm rectangle ROI prior and after application of a point bleach at its center. (B) Typical images of the ROI before (0 ms) and after (54, 199 and 990 ms) photobleaching. At each acquisition time, the ROI signal is averaged along the short axis into a one-dimensional profile, yielding a 2D chronograph of the FRAP experiment. (C) Example of chronograph obtained from GFP signal in a MDCK cell, and schematic of the three benchmarked FRAP methods.

For each acquisition, three FRAP repetitions were performed with a 20 s interval between repetitions to allow for complete fluorescence recovery.

#### 2.1.3 FRAP data pre-processing

The width of the rectangular ROI being nearly an order of magnitude smaller than effective bleaching radius, the recovery process is nearly invariant along the ROI width. Thus each image was averaged transversally resulting in a one dimensional time-dependent recovery profile.

In live-cell experiments, average background noise was estimated from extracellular regions and subtracted to the 1D recovery profile. Then, to account for fluorescence heterogeneity [25, 16] (Figure 1A&B), the 1D recovery profile after bleach was normalized by the 1D fluorescence profile before bleach. Finally, to account for fluorescence fading during the recovery phase, a boundary normalization was applied to the recovery profiles: the signal was normalized by the signal intensity at the extremities of the rectangular ROI (16% of the rectangle at each side) [14].

### 2.2 The impulse response (ImpRes) model for FRAP data analysis

Here we propose the derivation of a simple FRAP model to quantify the diffusivity of a free-diffusing probe based on a 1D recovery profiles obtained from the FRAP protocol described above.

We assume that the recovery is driven by purely Fickian diffusive behavior and that no transient binding, retention, or reaction processes are present. The transport of an inert species of concentration *C* with diffusivity *D* in an infinite homogeneous medium is described by a diffusion equation

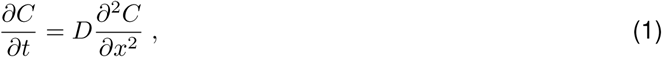

where *x* is the position and *t* is the time. The fundamental solution of this equation subject to an initial Dirac pulse at (*x, t*) = (0, 0) is a Gaussian function

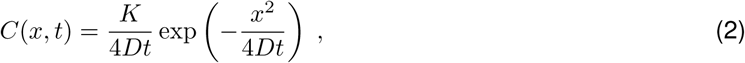

with *K* a multiplicative constant. Note that under the hypothesis above, this concentration profile remains valid for any time *t*, with constant values *K* and *D*.

In the case of FRAP experiments, this concentration corresponds to the bleached probe concentration profile. Therefore if the total (bleached and fluorescent) probe concentration *C_tot_* is constant, the normalized concentration of fluorescent probe *C^∗^* is just

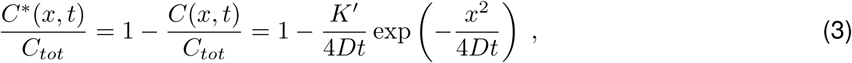

where *K^′^* = *K/C_tot_*is a constant. Finally, assuming the fluorescence intensity *I*(*x, t*) to be proportional to the fluorescent probe concentration, we simply have

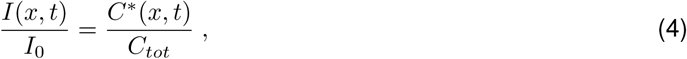

with *I*_0_ the fluorescence before bleaching (*ie* proportional to the total probe concentration).

In practice the initial bleaching phase is not a Dirac impulse but a spread Gaussian, thus fitting Equation (3) on experimental recovery profile is not straightforward. To circumvent this difficulty, we observe that everything occurs as if a Dirac impulse was set at a negative time, and that we can only observe the subsequent diffusion process for positive times. We thus define a virtual time delay *τ >* 0 after which a Dirac pulse would results in the Gaussian-like experimental observation. For a purely diffusive process, this concentration profile will remain Gaussian at any further time. Thus we can re-scale the time by defining *t^′^* = *t − τ*, so that the normalized fluorescence is

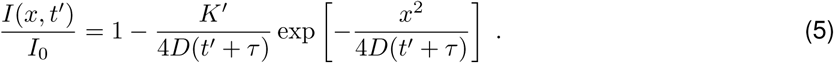

This equation is easily fitted to pre-processed experimental fluorescence profiles using the first postbleach acquisition time point as *t^′^* = 0. Three independent constants must then be determined: *K^′^* a constant proportional to the bleaching depth, *τ* the virtual time delay, and *D*, the probe diffusivity.

One advantage of this approach compared to most other methods is that it does not require any initial condition and thus does not give particular importance to the first post-bleach image. Moreover, this approach does not focus on a very narrow central ROI as do many existing methods. Instead, it uses all the information from the spatiotemporal fluorescence profile *I*(*x, t^′^*) to conduct a single least square fitting procedure with the three fitting parameters *K^′^*, *D*, and *τ* (Figure 1C).

### 2.3 FRAP datasets preparation

#### 2.3.1 *In silico* dataset: numerical generation of non-ideal FRAP recovery

To test the FRAP model in controlled settings, we performed numerical simulations of 1D FRAP recovery profiles, *I*(*x, t*), under both ideal and non-ideal conditions. Ideal recovery profiles were generated analytically by assuming an isotropic, homogeneous, and infinite medium in the in-plane directions. By assuming an initial Gaussian fluorescence profile that is invariant along the z-axis (out-of-plane), the solution to the diffusion equation is given by

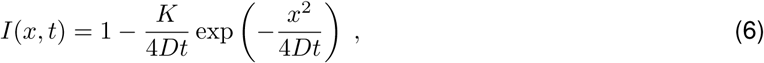

To mimic the sampling rate and spatial resolution of live-cell experiments, chronographs were constructed by sampling the analytical profiles at 20 ms temporal intervals and 0.1 µm spatial increments. The spatiotemporal domains were varied to assess model performance, with time ranges from 200 to 4000 ms and spatial extents from *±*12.5 to *±*50 µm.

To simulate realistic experimental noise, a stochastic term was added to Equation (6). This term was generated for each time step as a random 1D profile with a normal distribution. To ease visualization and mimic the finite resolution of the detector, the noise term was smoothed using a moving average filter over three time steps (60 ms) and three spatial positions (0.3 µm). To test the model under varied noise regimes, the noise standard deviation was tested at 0.150 (strong noise), 0.075 (medium noise), and 0.030 (weak noise).

Furthermore, the effect of imperfect boundary normalization was simulated by dividing the generated recovery profiles by the mean intensity of the first and last 40 pixels of the ROI. For non-ideal cases, this normalization was applied after the addition of the noise term, thereby introducing noise-induced artifacts into the normalization process, as it typically occurs in experimental data.

#### 2.3.2 *In vitro* dataset: Glycerol sample preparation

In vitro FRAP experiments were conducted using 20 kDa FITC-Dextrans (Sigma, fd20s 60842-46-8) immersed in a mixture made of autoclaved water and glycerol (Sigma, G2289-500ml, 3.3 nm hydrodynamic radius according to manufacturer data). FITC Dextran powder was added to hot water (60*^◦^*C) and stirred thoroughly. A controlled mass of glycerol was inserted in a Falcon and heated to 60*^◦^*C, then the same mass of FITC-Dextrans solution was added to the Falcon and stirred. To create a thin-film sample (1-10 µm thickness), several PDMS layers were piled on top of each other in a fluorodish (FD35-100, World Precision Instruments) containing 1 mL of the FITC-Dextrans mixture and were then compressed by the fluorodish upper part. Experiments were carried at room temperature. The theoretical diffusivity of 20 kDa Dextran was estimated for each experiment with Stokes-Einstein equation, based on the measured room temperature, mixture composition, and FD20 hydrodynamic radius.

#### 2.3.3 Live cell dataset: cell culture and sample preparation

MDCK type II cells (ATCC CRL-2936) were maintained in Dulbecco’s Modified Eagle Medium (DMEM) supplemented with 10% fetal bovine serum (Biowest), 1% penicillin-streptomycin (Gibco), red-phenol, GlutaMax, 4.5 g/L D-glucose, and pyruvate (Gibco). Cultures were incubated at 37*^◦^*C in a 5% CO2 atmosphere and passaged twice weekly using 0.05% Trypsin-EDTA (Gibco). Mycoplasma-free status was confirmed using the MycoSPY Master Mix (Biontex).

To express free-GFP, cells were transduced with MISSION TRC3 ORF GFP Lentivirus Control (Sigma) and subsequently sorted via Fluorescence-Activated Cell Sorting (FACS). High-fluorescence populations were selected for confocal Z-stack imaging and Fluorescence Recovery After Photobleaching (FRAP) measurements.

Live cell samples were prepared 48 to 72 hours prior to imaging. Cells were seeded onto fibronectin-coated fluorodishes (Sigma Aldrich, 1003624379). Cells were washed with 1X PBS and immersed in phenol-free DMEM supplemented with 4.5 g/L D-glucose, L-glutamine, 25 mM HEPES, sodium pyruvate (Gibco), 10% FBS, and 1% penicillin-streptomycin. Experiments were performed at 10–30% confluence to ensure the formation of small cellular islands. Large, spread cells located at the periphery of these islands were selected for analysis.

### 2.4 Benchmarked FRAP analysis methods

To evaluate the accuracy and robustness of the proposed ImpRes methodology (Equation (5)), we benchmarked its performance against two widely adopted FRAP analysis procedures for Gaussian bleaching profiles (Figure 1C).

#### 2.4.1 Orthogonal fit methodology

The first is the “orthogonal fit“ approach based on Kang et al. [16], which decouples the spatial and temporal components of the recovery. In this method, the initial bleach profile is first analyzed via a spatial fit *f_x_* to determine the effective bleach radius *R*_0_

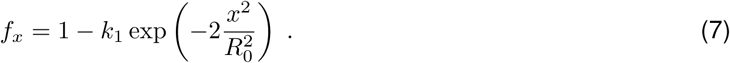

Subsequently, the temporal evolution of fluorescence within a central region of interest (ROI) with radius *R_ROI_* (set to 1 µm to match the bleach PSF) is analyzed via a temporal fit *f_t_*

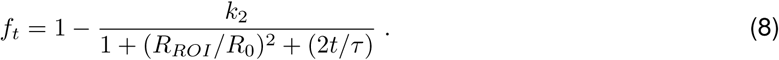

The resulting parameters, *R*_0_ and the recovery time *τ*, are then used to calculate the diffusion coefficient *D*

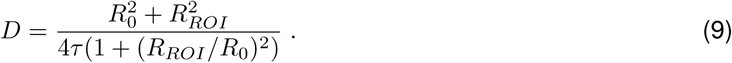

#### 2.4.2 Stacked fit methodology

The second is the “stacked fit” procedure based on Waharte et al. [24], which employs a sequential fitting strategy. At each time point *t_i_*, the spatial profile is fitted to a Gaussian function to determine the time-dependent squared radius *R*^2^(*t*)

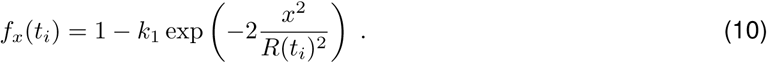

The resulting series of *R*^2^(*t*) values is then fitted temporally to extract the initial radius *R*_0_ and the diffusivity

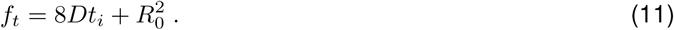

Similar to the ImpRes model, both the orthogonal and stacked methods were applied to our preprocessed 1D recovery profiles from live-cell experiments, in vitro standards, and numerical simulations. To rigorously assess the sensitivity of each model, we varied the fitting time windows by adjusting the initial delay (discarding early post-bleach frames) and the total fit duration (truncating late recovery frames). Representative fits for each methodology using simulated data are provided in Figures S8, S9, and S10.

## 3 Results

We aim to assess how optical biases can impact FRAP results and to what extent the proposed FRAP protocol and the ImpRes FRAP method could enhance robustness against (i) noise (ii) imperfect fluorescence fading correction by boundary normalization (iii) reduction of the fitted time-range.

We thus systematically compared ImpRes performance against two popular FRAP methods here referred to as the Orthogonal fitting method [16], and the Stacked fitting method [24] (see Figure 1C and Materials and Methods for details). We applied these methods to three FRAP datasets of increasing complexity obtained respectively *in silico*, *in vitro*, and in live cells.

### 3.1 *In silico* sensitivity analysis: ImpRes method is robust to noise and biased boundary normalization

To evaluate the accuracy and robustness of the proposed ImpRes methodology, we benchmarked it against the Orthogonal and Stacked fitting models using simulated Gaussian recovery profiles with a known probe diffusivity of *D_true_* = 20 µm^2^/s, a value representative of freely diffusing nanometric probes in the cytoplasm, such as GFP [26, 27, 28].

As a baseline, we first applied all three methods to ideal analytical Gaussian recovery profiles (Figure S1). As expected, all three methods accurately recovered the theoretical diffusivity across all tested fit delays and durations.

We next examined the sensitivity of these models to white noise, mimicking the low signal-to-noise ratios (SNR) often encountered in live-cell experiments with fast-diffusing probes (Figure 2A). We observed that the impact of noise on all methods is dependent on the fitting time range (Figure 2C). Because early postbleach images typically provide the deepest bleaching depth and best SNR, discarding them with a 200 ms fit delay dramatically increased the variance of the results across all methods and durations. The effects of increased fit duration varied by method. For the Stacked method, adding late recovery images did not improve accuracy; Gaussian fits performed on the low-SNR late images tended to destabilize the temporal fitting procedure (Figure S10), leading to significant inference biases. This suggests the Stacked method is only suitable for short analysis durations when dealing with noisy data.

**Figure 2:**
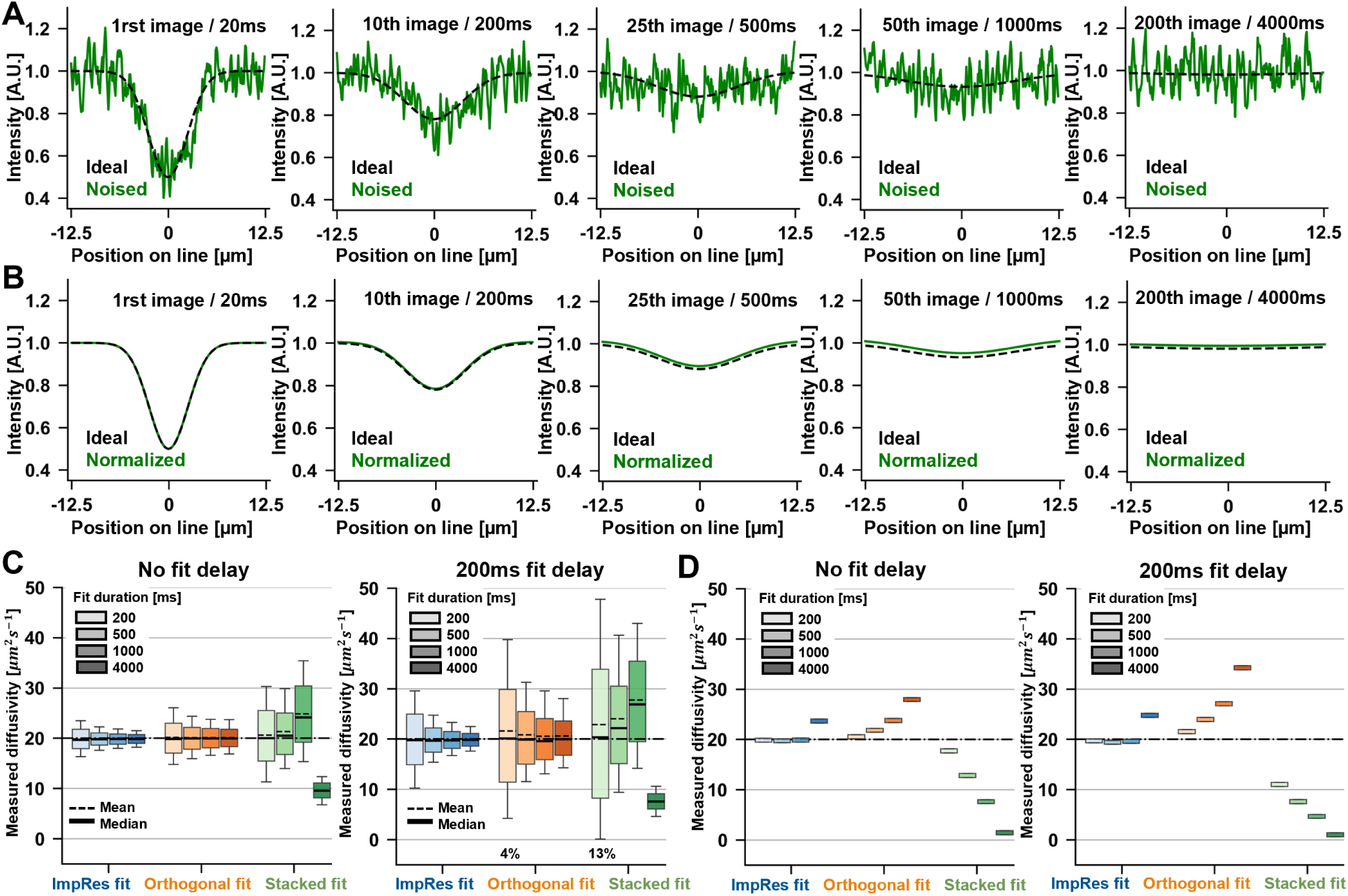
Sensitivity analysis of FRAP models to noise and boundary normalization using simulated signal. True diffusivity is 20 µm^2^/s. (A) Example of simulated ideal and noised Gaussian recovery profiles in the absence of boundary normalization. Noise standard deviation is 0.075. (B) Gaussian recovery profile with and without boundary normalization, in the absence of noise (C) Comparison of model estimation of diffusivity on simulated noised Gaussian recovery, without boundary normalization, for several fit durations. Noise standard deviation is 0.075. (Left) without fit delay, (right) with a 200 ms fit delay. Boxes are 1st and 3rd quartiles and whiskers are 10th and 90th percentiles. Percentages above and below boxes represent the proportion of points outside the fit threshold window [0.1 ; 100] µm^2^/s. Proportions <1% are not indicated. N=2000. (D) Comparison of model estimation of diffusivity on simulated Gaussian recovery with boundary normalisation, in the absence of noise, for several fit durations. (Left) without fit delay, (right) with a 200 ms fit delay. Thick colored bars are the obtained values, and do not represent inter-quartile range.

In contrast, both the ImpRes and Orthogonal methods gained precision as the fit duration increased from 200 to 4000 ms; the data spreading was reduced by half for ImpRes and by a third for the Orthogonal method. Notably, the spread was consistently lower for ImpRes than for the Orthogonal method by a factor of approximately 50%, depending on the time range. This is attributed to the fact that the Orthogonal method relies on a spatial fit from a single time step to determine the Gaussian radius *R*_0_ and subsequently the diffusivity, meaning that increasing the fit duration does not reduce the uncertainty associated with this parameter. Conversely, the ImpRes method utilizes all time points to optimize the spatial spreading, overcoming this limitation. Similar trends supporting the use of a single spatiotemporal fit were observed when the noise level was multiplied by 200% (Figure S6) or by 40% (Figure S5).

To mitigate low SNR, laser power is often increased during the recovery phase; however, this increases incidental photobleaching, which biases recovery images and requires correction. In live-cell experiments, utilizing a reference region is a common solution [14], provided the region is sufficiently distant to remain unaffected by the bleach. In practice, cellular dimensions often limit the distance of the reference region, and the boundaries may be slightly impacted by the spreading bleach profile. We therefore assessed the robustness of the three methods when using the longitudinal boundaries of the ROI (25 µm length) for signal normalization. As shown in Figure 2B, a slight discrepancy appears between the normalized and ideal Gaussian profiles at intermediate timescales. Surprisingly, applying the three methods to these normalized profiles (Figure 2D) revealed that this bias can dramatically impact the determination of the diffusivity, even in the absence of noise, particularly when early images are discarded or late time points are included. Both the Orthogonal and Stacked methods, which decouple time and space, were severely affected. The Orthogonal method deviated by up to 75% in the worst case (200 ms delay, 4000 ms duration), while the Stacked method appeared incompatible with this normalization except when limited to the first 200 ms without delay. While ImpRes was also affected by imperfect boundary normalization, the bias remained limited for most fit ranges and did not significantly compromise diffusivity inference.

To more closely simulate live-cell conditions, we analyzed the combined effect of noise and boundary normalization. We observed both a deviation from the expected diffusivity *D_true_* =20 µm^2^/s and an increase in result spreading across all models and time ranges (Figure 3A). Comparison with previous results shows that normalizing noisy data exacerbates the variance, particularly when very late recovery images are included. However, the normalization bias itself was not strongly influenced by noise. These results confirm that the ImpRes method systematically yields less biased results with lower variance.

**Figure 3:**
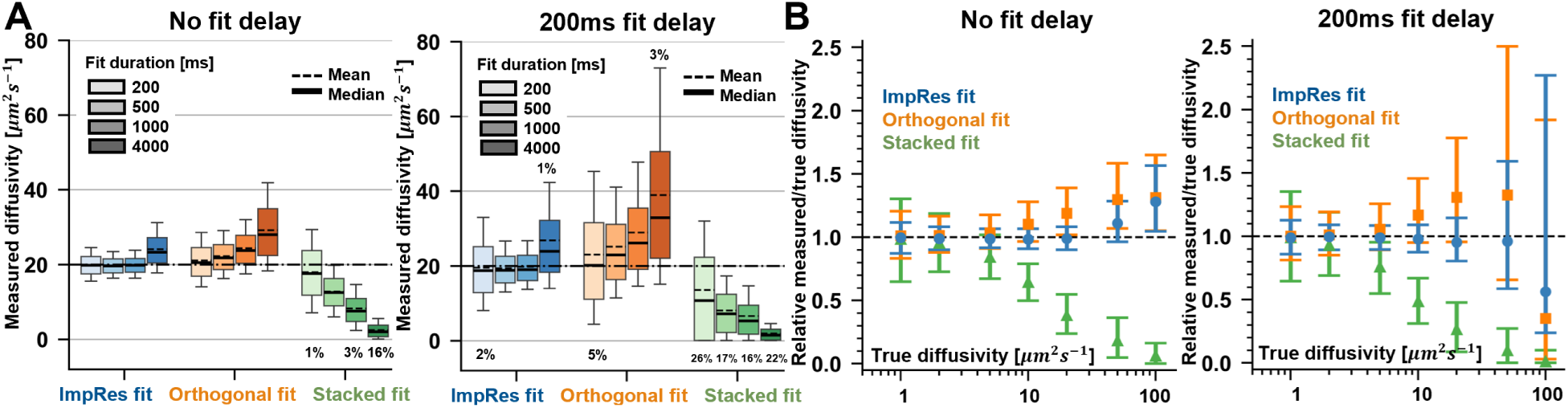
Comparison of the three FRAP models for an intermediate fit duration (1000 ms), with and without fit delay (200 ms) for simulated Gaussian recovery profiles with boundary normalization. True diffusivity is 20 µm^2^/s, noise standard deviation is 0.075, N=2000. (A) Measured diffusivity obtained without fit delay (left) and with a 200 ms fit delay, for various fit durations. Boxes are 1st and 3rd quartiles and whiskers are 10th and 90th percentiles. Percentages above and below boxes represent the proportion of points outside the fit thresholds window [0.1 ; 100] µm^2^/s. Proportions <1% are not indicated. (B) Relative measured/true diffusivity as a function of true diffusivity, without fit delay (left) and with 200ms fit delay (right), for true diffusivity ranging from 1 to 100 µm^2^/s. Errorbars are 1st and 3rd quartiles, point is median value.

Finally, to determine the applicable diffusivity range for each method, we studied the combined effect of noise and normalization for *D_true_*ranging from 1 to 100 µm^2^/s at an intermediate fit duration of 1000 ms (Figure 3B). All methods performed well for low diffusivities (*D_true_ ≤* 2 µm^2^/s), although the variance was highest for the Stacked method and lowest for ImpRes. For higher diffusivities, only the ImpRes method maintained an error *≤*10% up to *D_true_* =50 µm^2^/s, even with a 200 ms fit delay. Although the accurate determination range for each method can be tuned by adapting the fit duration to the recovery kinetics (Figure S7), the ImpRes method consistently provided a wider domain of validity and lower variance.

### 3.2 *In vitro* validation: ImpRes method is insensitive to ROI length

To further validate the proposed methodology and confirm the findings from our numerical simulations, we performed in vitro FRAP experiments using FITC-Dextran (20 kDa) in a glycerol-water mixture (50% by mass). Depending on the precise ambient temperature, the expected diffusivity for this system is approximately 10 to 20 µm^2^/s, which mimics the diffusion coefficients of intracellular probes such as free-GFP.

Given that fluorescence fading correction is a critical step in FRAP analysis, and that utilizing a reference region too close to the bleached area can introduce significant bias – particularly for the Orthogonal and Stacked methods – we evaluated the impact of ROI geometry. Experiments were conducted in thin glycerol films (1–5 µm thickness) using a rectangular ROI of 30 µm, which was then numerically cropped to study the influence of the ROI lengths on the models performances.For the full 30 µm ROI without fit delay, all three methods recovered a consistent diffusivity of about 12 µm^2^/s (Figure 4A), which is in good agreement with the theoretical value derived from the Stokes-Einstein equation (11 µm^2^/s). The minor discrepancy is likely due to slight variations in the glycerol-water mass fraction and ambient temperature. These results indicate that all three methods are applicable to experimental data obtained with the proposed FRAP protocol provided the ROI is sufficiently long, thereby validating the experimental setup. However, as the rectangle length was gradually reduced, the Stacked method failed to converge and the Orthogonal method exhibited significant bias. In contrast, the diffusivity obtained by the ImpRes method remained largely unaffected.

**Figure 4:**
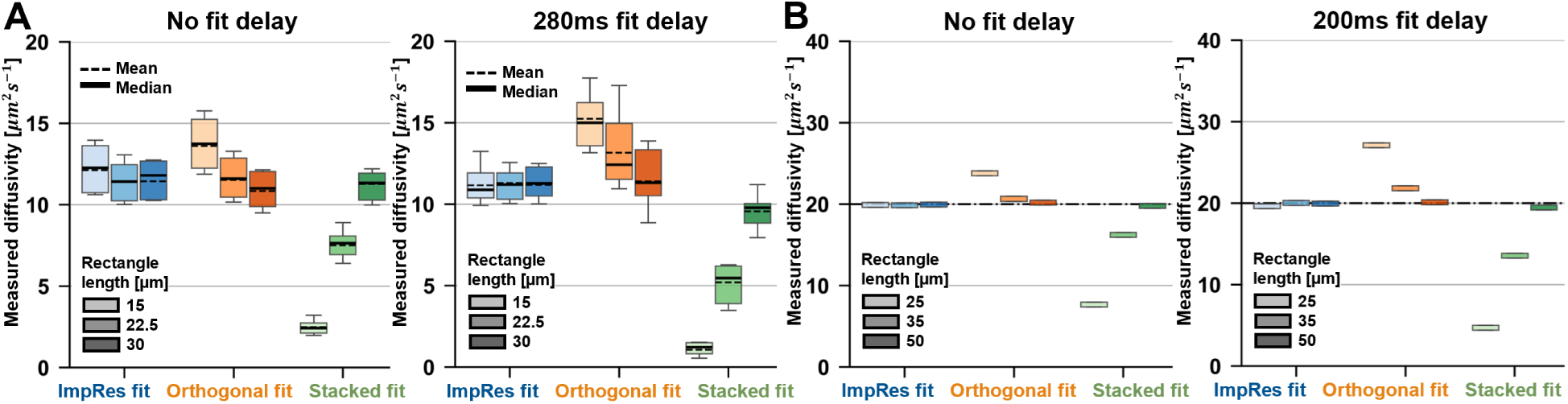
Influence of rectangular ROI length on FRAP models robustness water-glycerol mixtures. (A) Diffusivity measured in 20 kDa FITC-Dextrans in 50/50 (by weight) glycerol-water at 24*^◦^*C, either without fit delay (left) or with (right) a 280 ms fit delay, for several rectangular ROI lengths. Expected diffusivity is about 11 µm^2^/s. Boxes represent 1st and 3rd quartiles and whiskers are 10th and 90th percentiles. Two experiments were conducted for a total of 10 measurements per condition. (B) Numerical results obtained on Gaussian recovery profile with boundary normalization, either without (left) or with (right) a 200 ms fit delay, for several rectangle lengths. Thick colored bars are the obtained values, and do not represent inter-quartile range. True diffusivity is 20 µm^2^/s.

To determine if these observations were consistent with our previous numerical findings, we performed simulations of non-noisy, normalized diffusion profiles for various rectangle lengths at *D_true_*=20 µm^2^/s (Figure 4B). The resulting trends for the three models are qualitatively consistent with the experimental data from the glycerol mixture (Figure 4A). This comparison not only validates our numerical results but also suggests that increasing the fit duration and reducing the ROI length have qualitatively similar effects on normalization-induced bias. Specifically, the robustness of the ImpRes method to longer fit durations appears to be directly linked to its lower sensitivity to reduced ROI lengths, favoring its use inside finite-size living cells.

### 3.3 Live cell application: ImpRes method gives consistent diffusivity even in the presence of short-time artifacts

Finally, the proposed FRAP protocol and the ImpRes analysis framework were applied to MDCK cells expressing cytoplasmic free-GFP. Based on the expected diffusivity of about 20 µm^2^/s and our previous benchmarking results, recovery was imaged using a 25*×*1 µm rectangular ROI at a frequency of 50 Hz.

Pre-processed data from 21 cells were analyzed using the three benchmarked models across various fit durations and delays (Figure 5). We observed that all methods exhibited some degree of sensitivity to the fitting time range, suggesting the presence of a systematic bias. Specifically, the average recovery profile revealed an unexpected transient increase in fluorescence during the early post-bleach phase, which gradually dissipated after 100–200 ms (Figure S2), leaving a nearly ideal Gaussian profile at later time points.

**Figure 5:**
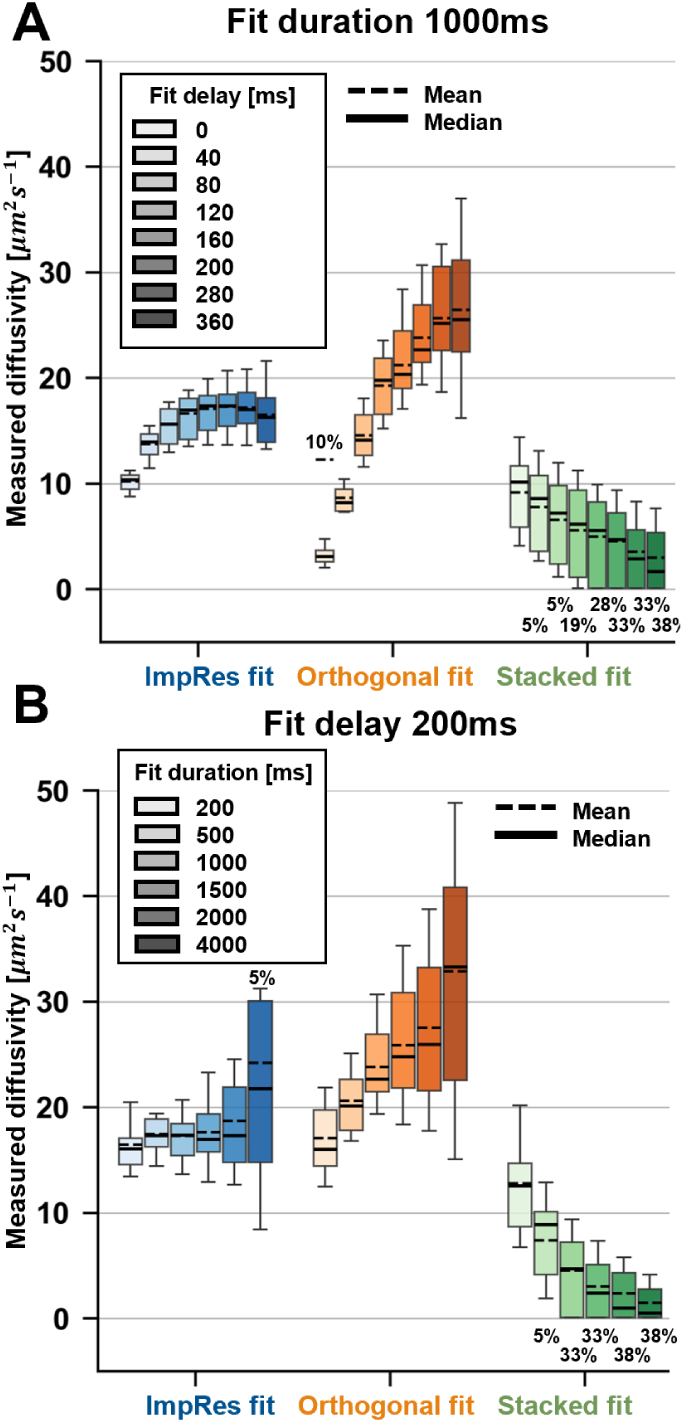
Free-GFP diffusivity obtained from the three FRAP models in MDCK cells cytoplasm. (A) Diffusivity obtained for a 1000 ms fit duration as a function of fit delay. (B) Diffusivity obtained for a fit delay of 200 ms as a function of fit duration. Boxes are 1st and 3rd quartiles and whiskers are 10th and 90th percentiles. Percentages above and below boxes represent the proportion of points outside the fit thresholds window [0.1 ; 100] µm^2^/s. Proportions <1% are not indicated. Results obtained on 21 cells. photobleaching, but sufficient energy to trigger a transient increase in fluorescence. The subsequent disappearance of this population after about 100 ms could be explained by the delayed de-excitation of these probes.

Consistent with this observation, the ImpRes model results stabilized for fit delays above 100 ms, supporting the reliability of a fit-delay window between 80 and 360 ms (Figure 5B). When a 200 ms fit delay was applied (Figure 5A), the ImpRes results became largely independent of the total fit duration, yielding an estimated diffusivity of approximately 18 µm^2^/s. This robustness was not observed for the Stacked or Orthogonal methods. Based on our previous results (Figures 2 and 3), we attribute this sensitivity to the limited robustness of the Stacked and Orthogonal models against imperfect boundary normalization. Additionally, the ImpRes method demonstrated lower sensitivity to noise, as evidenced by its reduced variability compared to the Orthogonal method.

To determine the origin of the observed transient fluorescence increase, we first investigated whether 3D diffusion effects in MDCK cells contributed to the bias. The assumption of axial invariance is often compromised by sharp variations in cell height or the use of high-numerical-aperture objectives [18]. To test this possibility, we performed supplementary experiments using FITC-Dextran in glycerol-water mixtures with sample heights varying from 2 to 9 µm. No transient rise in fluorescence was observed; instead, increasing sample thickness merely resulted in an enlargement of the bleached profile (Figure S3), consistent with the hourglass shape of the bleaching PSF [18]. Furthermore, while sample thickness did influence the inferred diffusivity (Figure S4), the effect was significantly smaller than the bias observed in the live-cell GFP data (Figure 5).

We therefore hypothesize that the bias originates from the complex photophysical properties of GFP. Various GFP variants in specific cell lines have been reported to exhibit flickering, intermediate fluorescence states [20, 21], photoactivation [22, 29, 30], or photoswitching [31]. We propose that during the bleaching phase, GFP molecules adjacent to the focal center receive an insufficient intensity for complete

## 4 Discussion

In this work, we proposed and validated a robust yet simple FRAP methodology designed for the quantification of cytoplasmic molecular diffusivity using standard laser scanning confocal microscopy (LSCM). The underlying model assumes a Gaussian signal recovery from an impulse response (ImpRes) enabling a single-step spatiotemporal fitting procedure that maximizes the information extracted from the recovery dataset while minimizing the influence of common experimental artifacts such as noise, biased boundary normalization, and complex probe photodynamics at early time. The performance of this model was systematically confronted to two state of the art methods with comparable simplicity used to analyse Gaussian bleaching profiles, here referred to as the Orthogonal method [16], and the Stacked method [24]. Applications to datasets of increasing complexity obtained *in silico*, *in vitro*, and in live cells, showed that the ImpRes method is consistently more accurate and precise in estimating diffusivities across a biologically relevant range. It is also more robust to poor signal to noise ratio, biased fluorescence normalization, unusable early postbleach images, and reduced ROI length.

While the Stacked fit approach also utilizes all recovery data, it remains sensitive to noise and normalization errors. We propose that the enhanced accuracy of ImpRes stems from its reliance on a single global fit of the impulse response of the diffusion equation. This approach enforces spatial and temporal consistency across the entire acquisition, effectively averaging out stochastic noise and mitigating the bias introduced by imperfect boundary normalization. This robustness is particularly critical for intracellular measurements, where the finite size of the cell often prevents the definition of a truly independent reference region, forcing a reliance on the ROI boundaries for normalization. The ImpRes method is also particularly robust to the ROI length, which facilitates the study of diffusion phenomena in confined environment. For even smaller compartments, the approach could be extended to shorter rectangle ROI by bleaching one extremity and using the other as a the fluorescence reference.

Our application of this protocol to turbo-GFP in live MDCK cells revealed an unexpected transient fluorescence rebound during the early post-bleach phase. Through a series of controlled in vitro experiments using FITC-Dextran in glycerol-water mixtures of varying thicknesses (2–9 µm), we observed that increased sample thickness can lead to a modest overestimation of diffusivity (approximately 10–30%) likely due to the hourglass-shaped bleach PSF characteristic of high numerical aperture objectives [18]. However this effect was insufficient to explain the magnitude of the rebound seen in live cells. Instead, these results suggest that the bias originates from the complex photophysical properties of the GFP variant. Given that GFP molecules can exhibit flickering, photoactivation, or photoswitching [20, 21, 22], it is probable that molecules at the periphery of the bleach zone undergo a transient increase in fluorescence due to suboptimal bleaching intensities. Because the ImpRes method is robust to the exclusion of early time points, we were able to recover a stable diffusivity of about 18 µm^2^/s. This value is slightly lower than those previously reported for monomeric GFP [27, 25], a discrepancy that may be explained by the tendency of turbo-GFP to form dimers, thereby increasing its hydrodynamic radius and reducing its diffusion coefficient [32, 33].

While this study focused on freely diffusing probes, the ImpRes framework can be readily extended to systems with a significant immobile or bound fraction. By utilizing late-stage recovery images to define the bound baseline, the mobile fraction can be recovered from experimental acqusition and then analyzed using the same spatiotemporal fit (Equation (5)). However, as with most FRAP-based methods, the inference of dynamics remains challenging in the presence of multiple diffusing populations or complex binding/unbinding kinetics that occur on the same timescale as diffusion. Importantly, even in cases where the ImpRes approach might be difficult to apply, our thorough investigation of the different biases impacting FRAP application in living cells may favor the correct implementation of other methods by emphasising the importance of signal to noise ratio, the reference region used for fluorescence fading correction, cell thickness, and other short-time artifacts.

In conclusion, the ImpRes approach offers a robust and computationally simple alternative to existing FRAP analysis models. While complementary techniques such as Single Particle Tracking (SPT) and Fluorescence Correlation Spectroscopy (FCS) provide deeper insights into the nature of intracellular heterogeneity and sub-diffusive motion [34], FRAP remains an indispensable tool due to its versatility and ease of implementation [10]. By investigating FRAP sensitivity to noise, boundary artifacts, and probe-specific photodynamics, and proposing a robust ImpRes framework against all these biases, this work expands the utility of FRAP for the quantitative study of fast-diffusing macromolecules in the complex environments of living cells.

## Code availability

All codes and example datasets are available on github.com/destriano/frap_imp_res.

## Acknowledgements

We are thankful to Benoît Ladoux for his insights and support along this project. We also thank Xavier Baudin from Imagoseine platform for helpful discussions on GFP complex fluorescence properties. We acknowledge the ImagoSeine core facility of the Institut Jacques Monod, member of the France BioImaging infrastructure (ANR-24-INBS-0005 FBI BIOGEN) and GIS-IBiSA. This work was supported by the LABEX Who Am I? (ANR-11-LABX-0071 to R.-M.M.), the Ligue Contre le Cancer (Equipe labellisée 2019 to R.M.M.) and the Agence Nationale de la Recherche (“STRATEPI” DFG-ANR-22-CE92-0048, and “VISCOMAG2” ANR-24-CE42-6142 to R.-M.M.).

## Supporting figures

**Figure S1:**
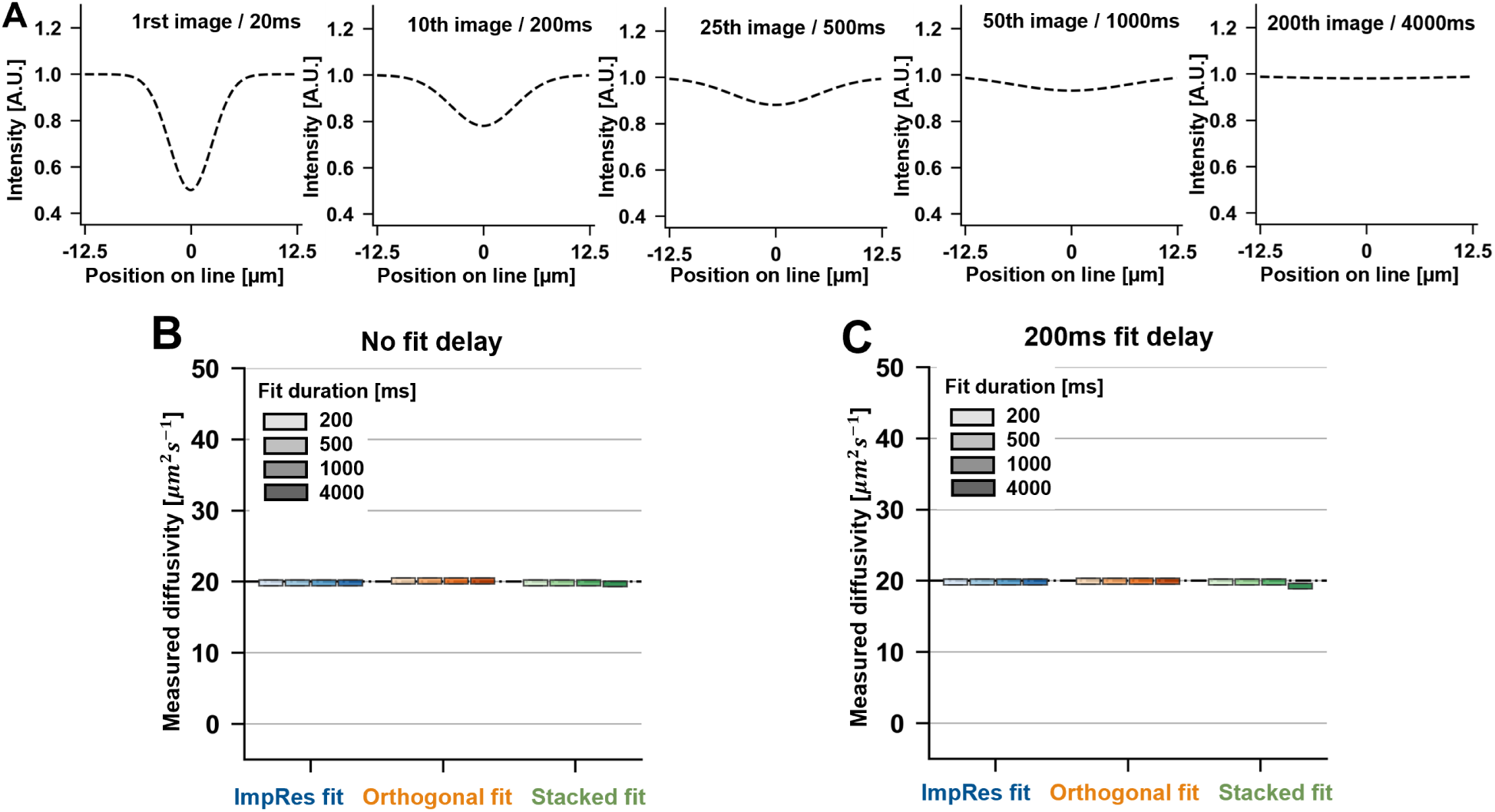
Application of the three benchmarked protocols to ideal Gaussian recovery curves for several fit duration and fit delays. All models recover the correct diffusivity, validating the implementation of the models and their applicability to Gaussian bleaching profiles. (A) Analytical Gaussian recovery profile for a 20 µm^2^/s diffusivity. (B) Results obtained without fit delay. (C) Results obtained for a 200 ms fit delay.

**Figure S2:**
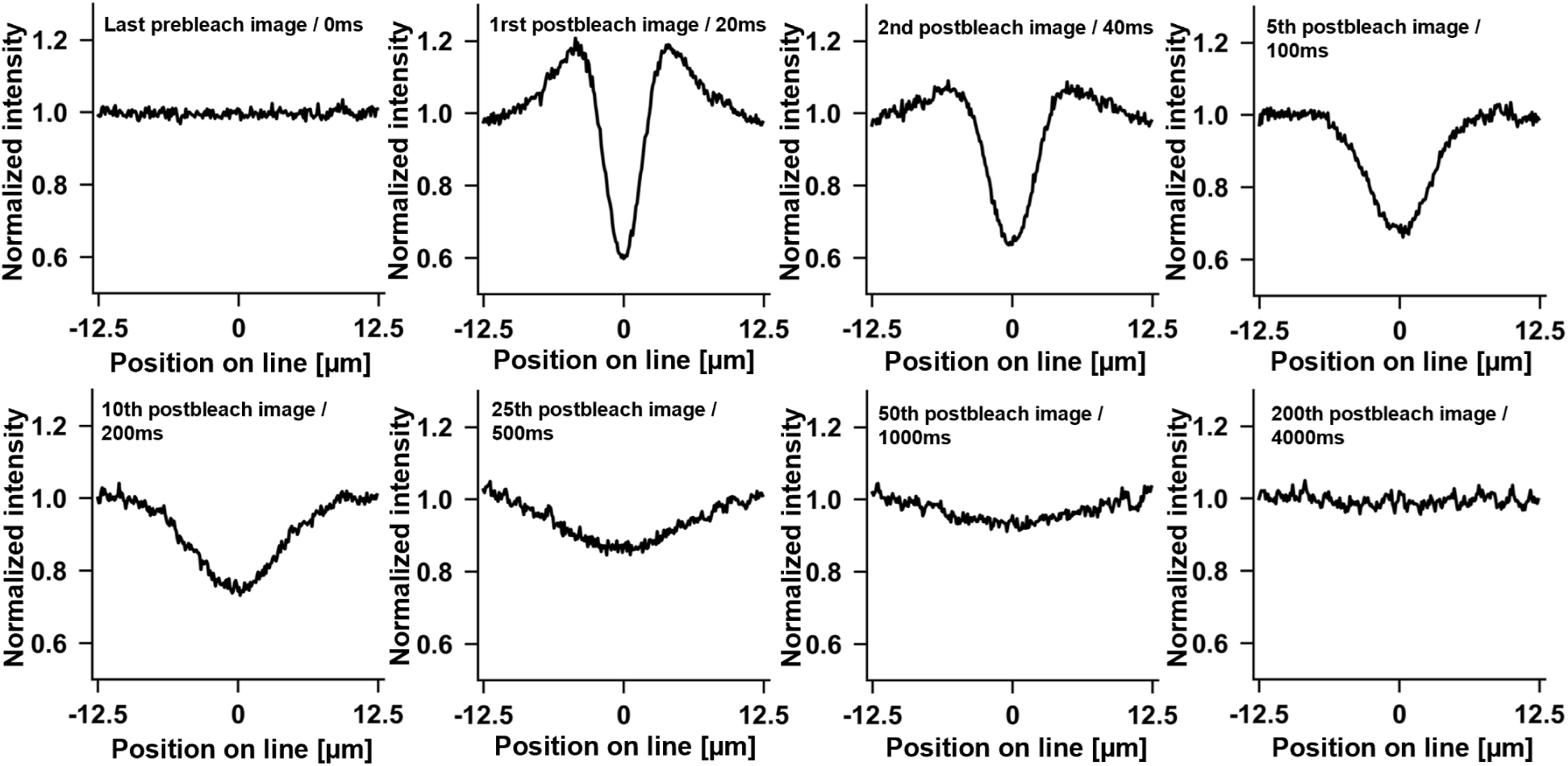
Average free-GFP fluorescence profile obtained by FRAP in 21 MDCK cells, with double normalization by prebleach profile and by rectangle boundaries. A strong deviation from Gaussian profiles is visible in the early postbleach images (<100 ms), making them unsuited for application of most FRAP diffusivity inference models. The bleaching profile becomes Gaussian after a few images, supporting the relevance of applying a fit delay for FRAP application inside living cells.

**Figure S3:**
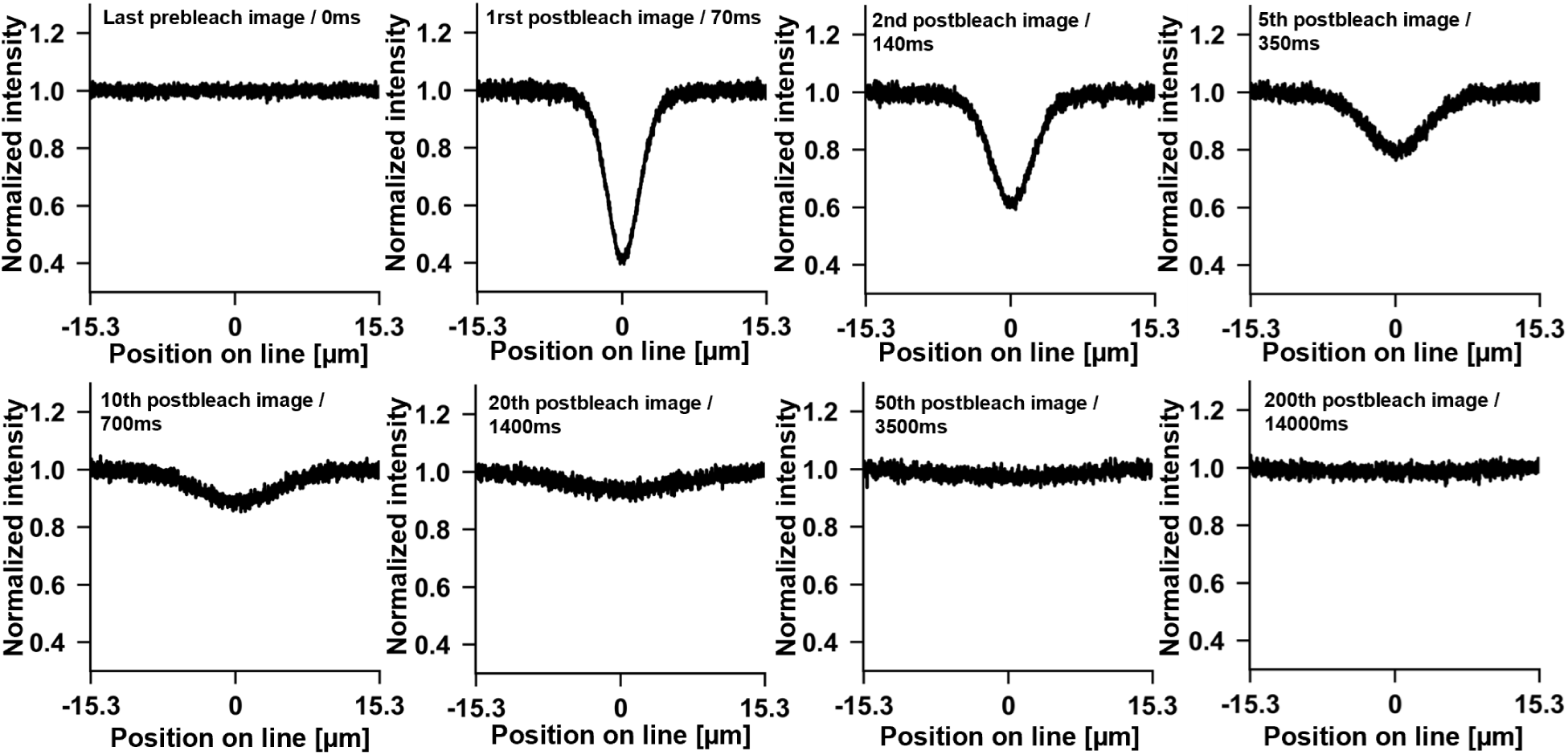
Average *in vitro* FITC-Dextran fluorescence profile obtained with FRAP protocol from 10 measurements in 2 independent experiments (measured diffusivity around ^1^2*µm*^2^*s^−^*^1^), with double normalization by prebleach profile and by rectangle boundaries. No strong deviation from Gaussian profile is visible at any timescale, the bleach profile stays Gaussian during the full recovery.

**Figure S4:**
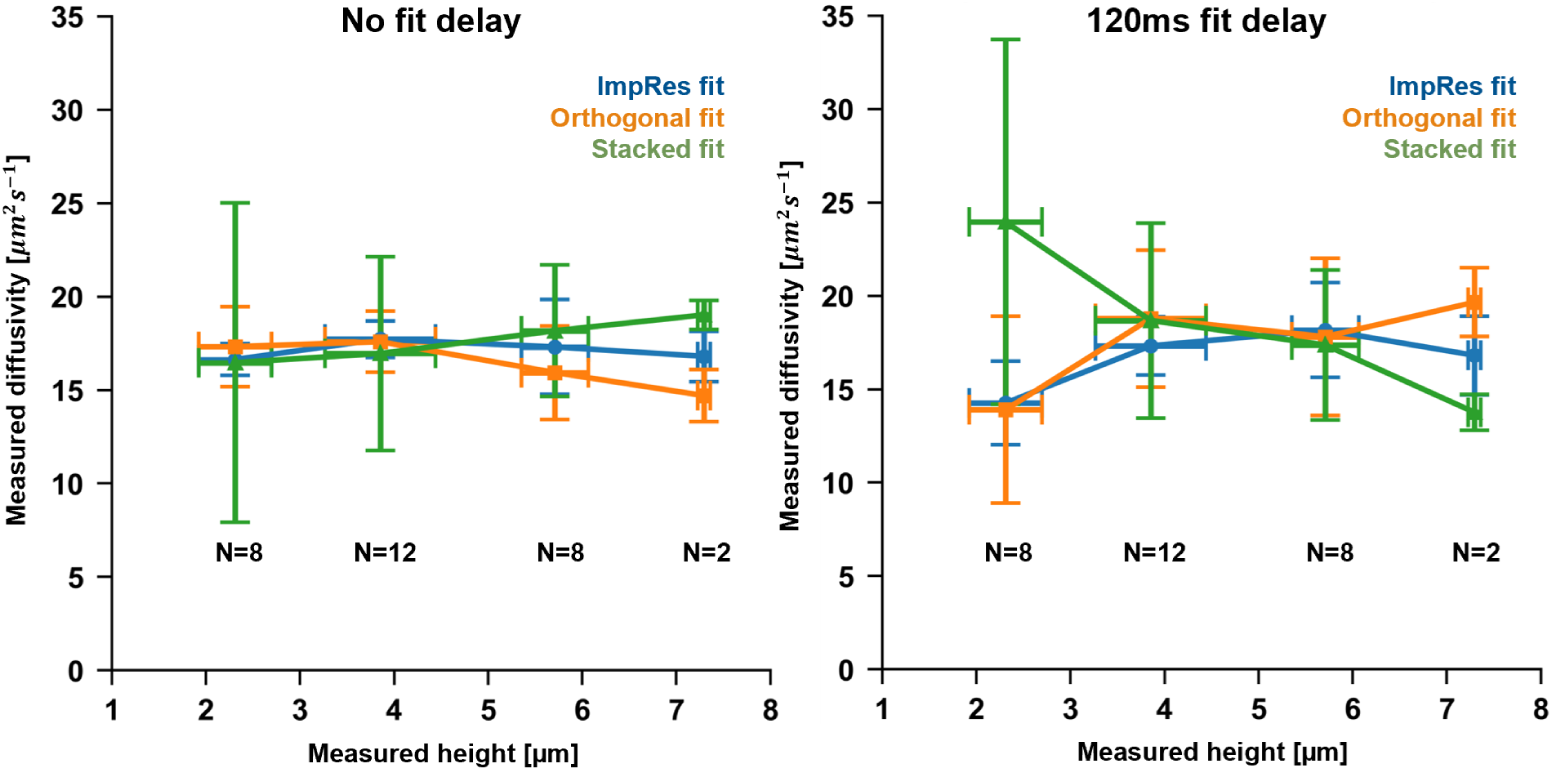
Impact of glycerol sample height on diffusivity inference using the three benchmarked FRAP methods on 20 kDa FITC-Dextrans in 47.5%/52.5% glycerol-water at 24*^◦^*C, either without (A) or with (B) a 120 ms fit delay. Expected diffusivity is 15 µm^2^/s. Errorbars are mean *±*std. Data was binned in [1,3], [3,5], [5, 7], [7, 9] µm height intervals, with corresponding number of measurements indicated on graph.

**Figure S5:**
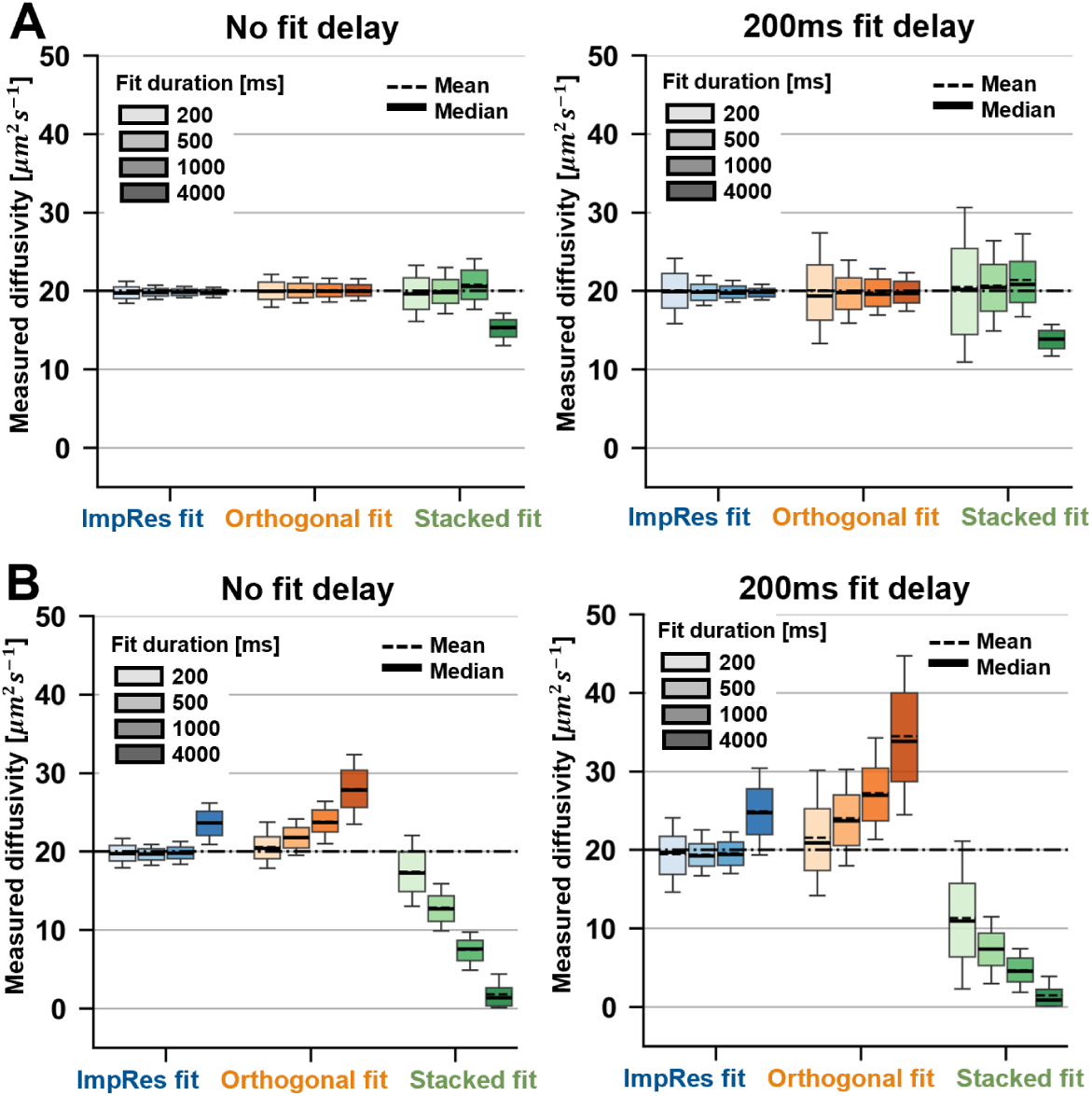
Diffusivity measured in weakly noised Gaussian recovery profiles with the three benchmarked FRAP models. Noise standard deviation is 0.030 (40% of the base level shown in main manuscript figures). (A) Without fit delay (left) and with a 200 ms fit delay (right), without profile normalization by its boundaries. (B) Without fit delay (left) and with a 200 ms fit delay (right), with profile normalization by its boundaries. N=400, boxes are 1st and 3rd quartiles and whiskers are 10th and 90th percentiles. True diffusivity is 20 µm^2^/s. When present, percentages above (respectively below) boxes represent the proportion of fits that reached the upper fit boundary at 100 µm^2^/s (respectively lower fit boundary at 0.1 µm^2^/s). When proportion is <1%, percentage is not indicated.

**Figure S6:**
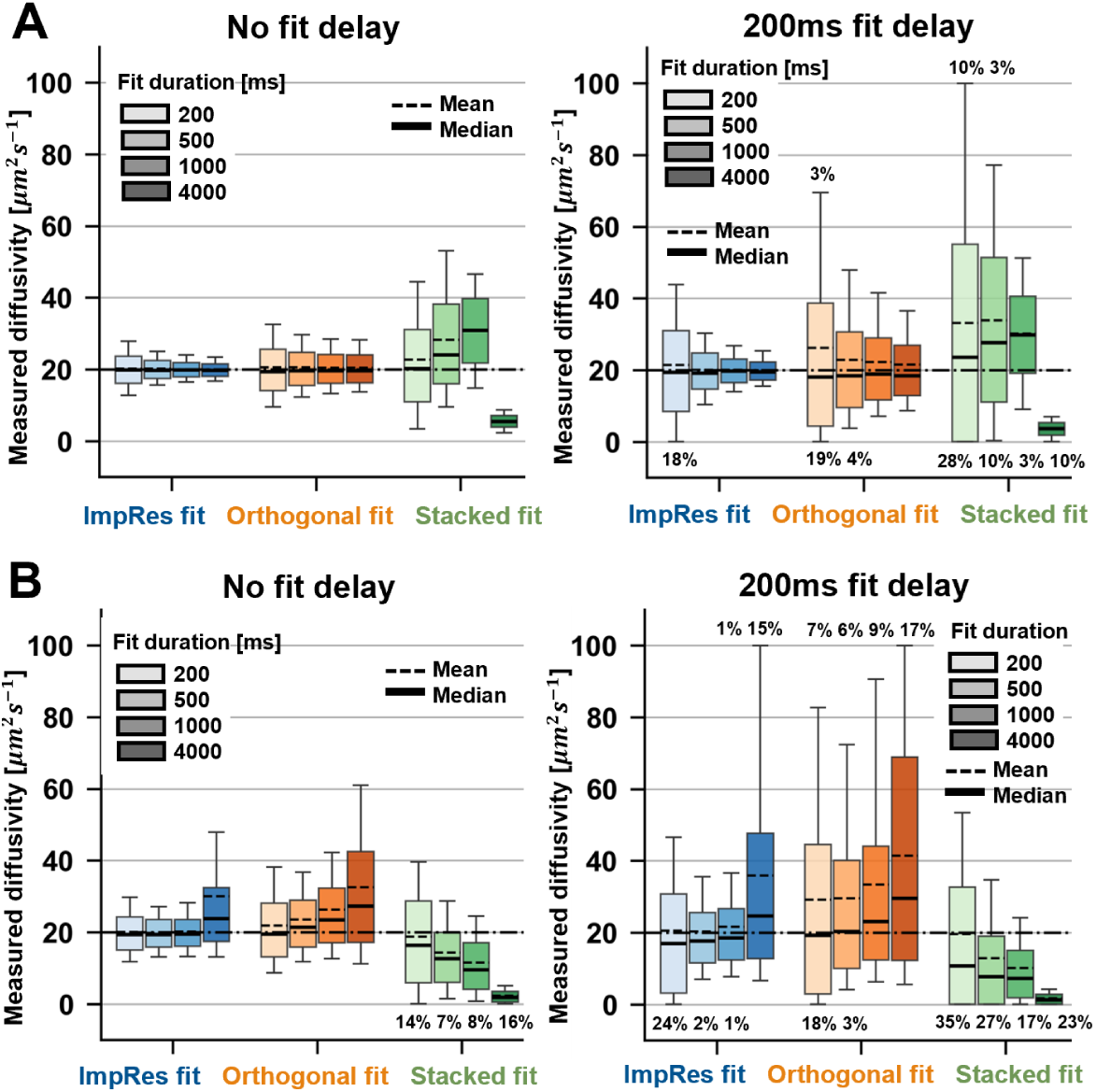
Diffusivity measured in highly noised Gaussian recovery profile with the three benchmarked FRAP models. Noise standard deviation is 0.150 (200% of the base level shown in main manuscript figures). (A) Without fit delay (left) and with a 200 ms fit delay (right), without profile normalization by its boundaries. (B) Without fit delay (left) and with a 200 ms fit delay (right), with profile normalization by its boundaries. N=1100, boxes are 1st and 3rd quartiles and whiskers are 10th and 90th percentiles. True diffusivity is 20 µm^2^/s. When present, percentages above (respectively below) boxes represent the proportion of fits that reached the upper fit boundary at 100 µm^2^/s (respectively lower fit boundary at 0.1 µm^2^/s). When proportion is <1%, percentage is not indicated.

**Figure S7:**
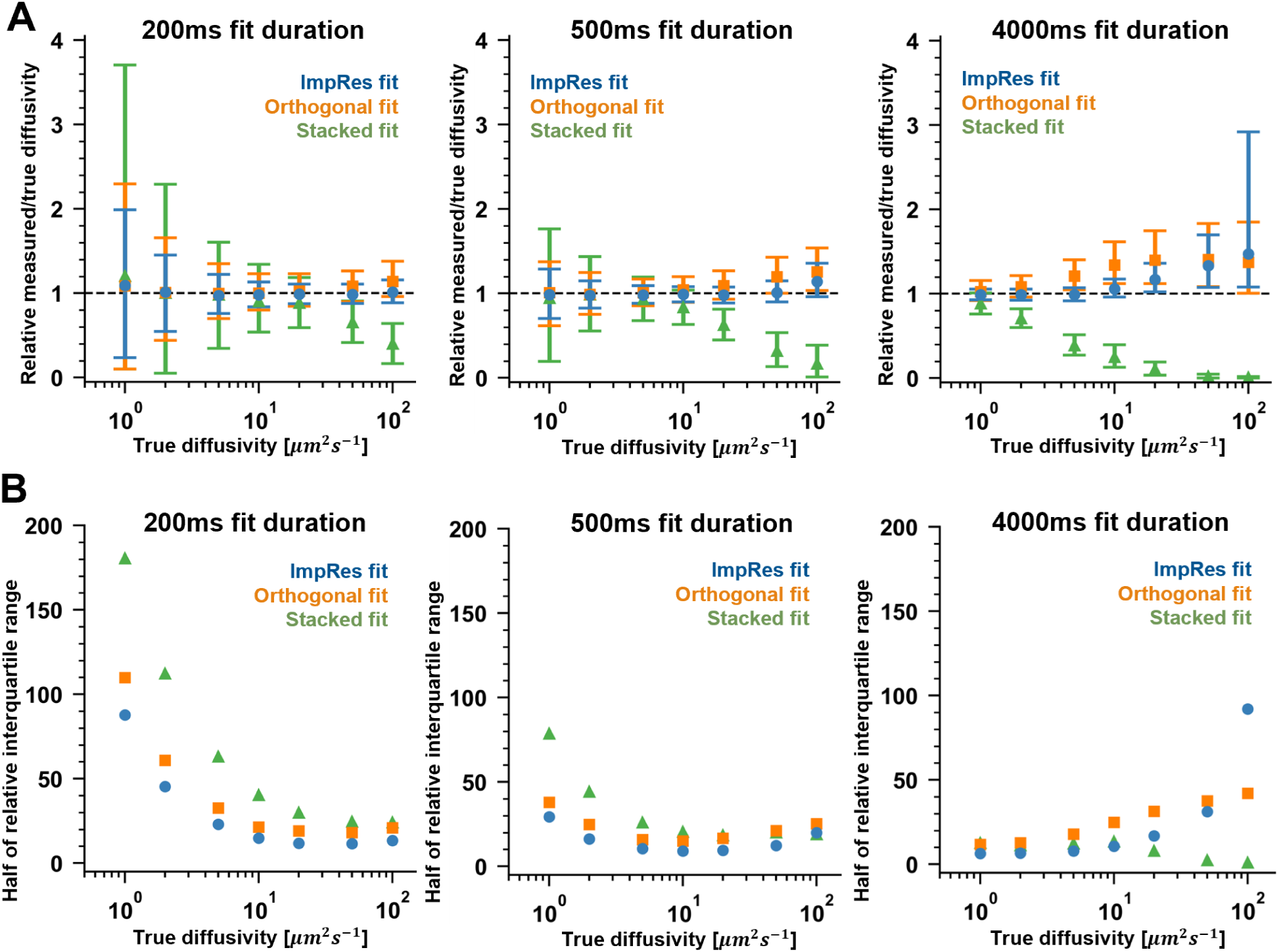
Comparison of the three FRAP models for several fit durations (200 ms left, 500 ms middle, 4000 ms right), without fit delay, for noised and normalized Gaussian recovery profiles (N=2000), for true diffusivity ranging from 1 to 100 µm^2^/s. Errorbars are Q1 and Q3, point is median value. (A) Measured diffusivity relative to true diffusivity. (B) Dispersion of the results quantified by half of the interquartile range relative to true diffusivity.

**Figure S8:**
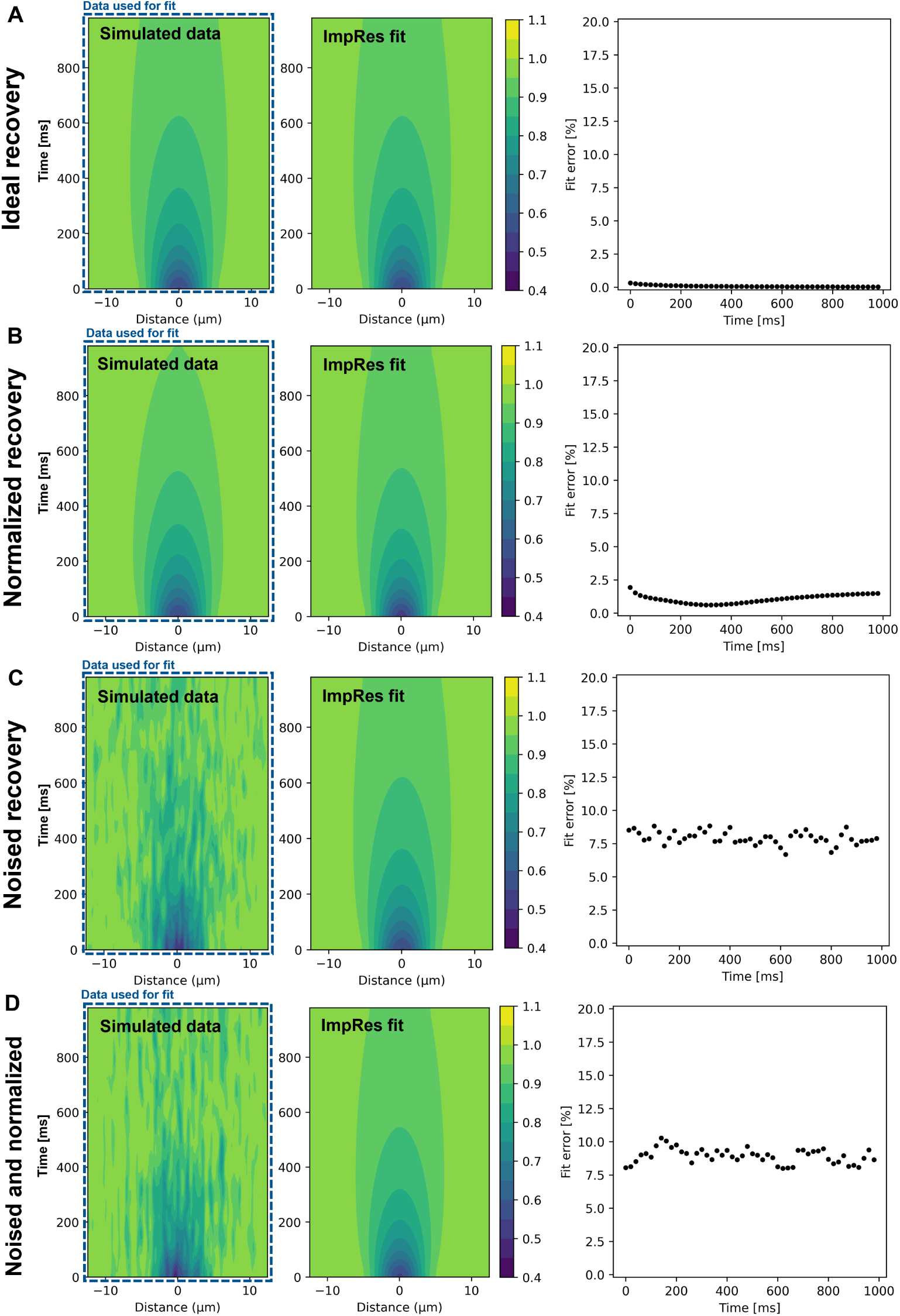
Examples of simulated data (*D*=20 µm^2^/s) and fit with ImpRes method for (A) ideal Gaussian recovery; (B) normalized recovery; (C) noised recovery, with noise standard deviation at 0.075; (D) Normalized and noised recovery, with noise standard deviation at 0.075.

**Figure S9:**
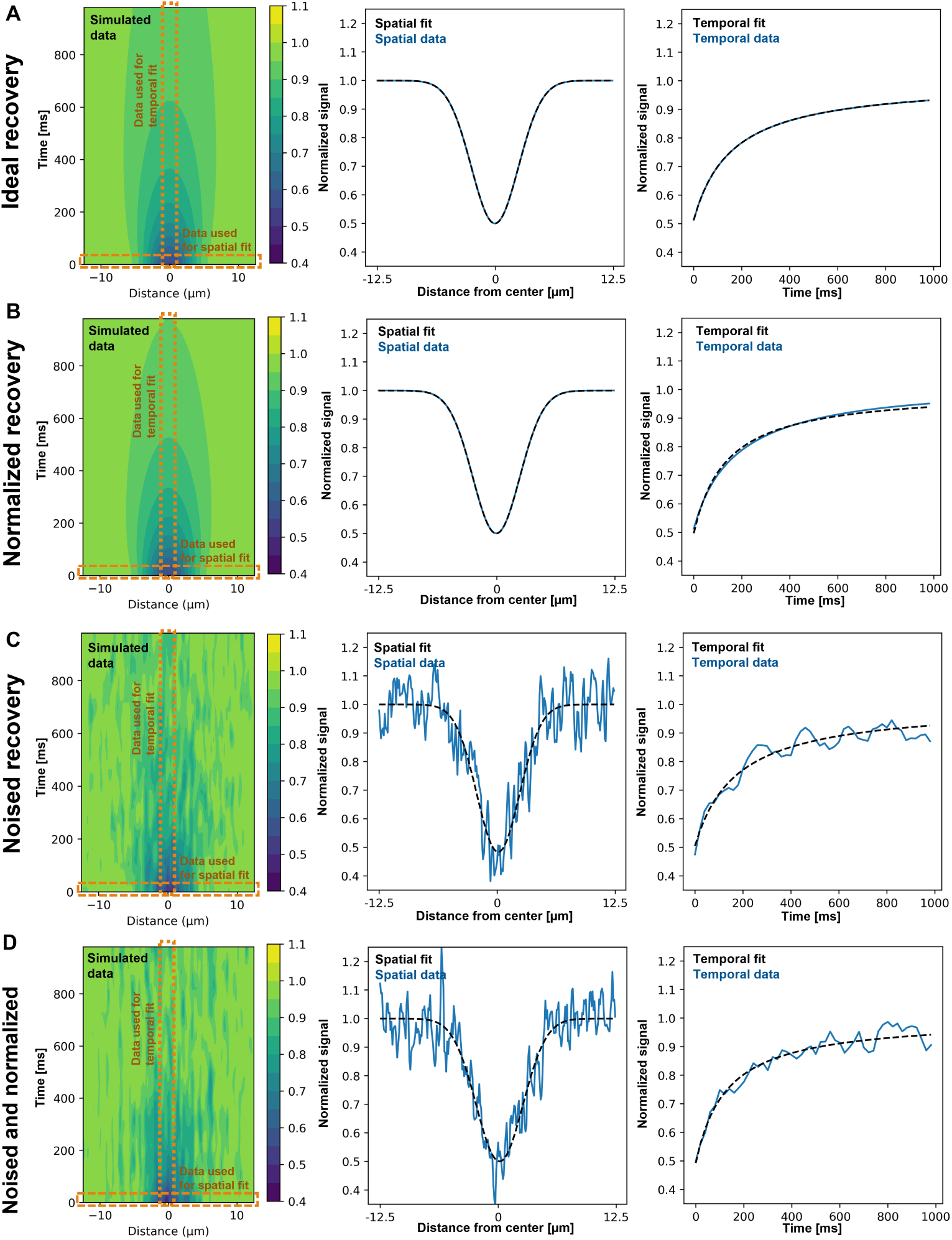
Examples of simulated data (D=20 µm^2^/s) and fit with Orthogonal method for (A) ideal Gaussian recovery; (B) normalized recovery; (C) noised recovery, with noise standard deviation at 0.075; (D) normalized and noised recovery, with noise standard deviation at 0.075.

**Figure S10:**
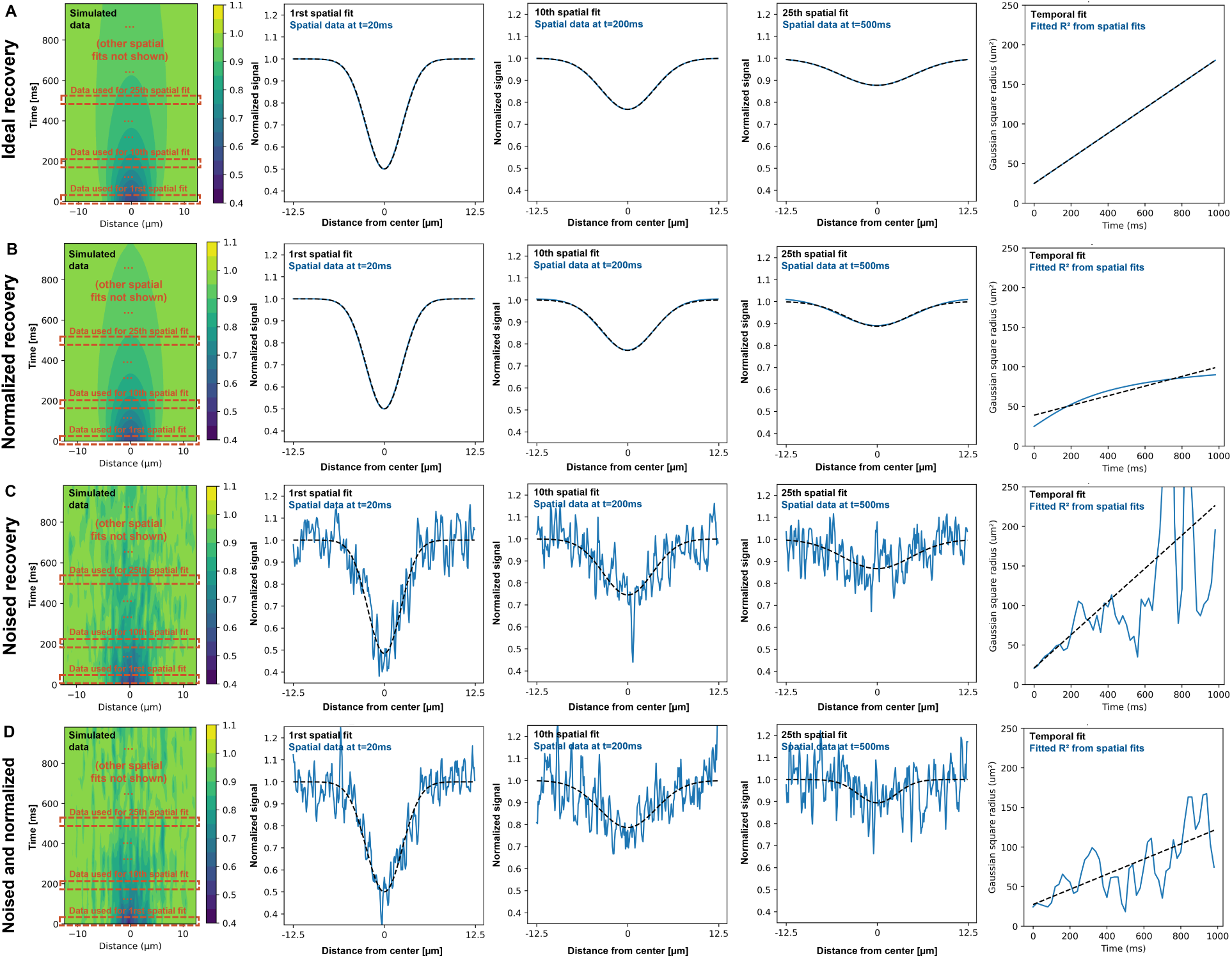
Examples of simulated data (*D*=20 µm^2^/s) and fit with Stacked method for (A) ideal Gaussian recovery; (B) normalized recovery; (C) noised recovery, with noise standard deviation at 0.075; (D) normalized and noised recovery, with noise standard deviation at 0.075.

